# Principles governing A-to-I RNA editing in the breast cancer transcriptome

**DOI:** 10.1101/012849

**Authors:** Debora Fumagalli, David Gacquer, Françoise Rothé, Anne Lefort, Frederick Libert, David Brown, Naima Kheddoumi, Adam Shlien, Tomasz Konopka, Roberto Salgado, Denis Larsimont, Kornelia Polyak, Karen Willard-Gallo, Christine Desmedt, Martine Piccart, Marc Abramowicz, Peter J Campbell, Christos Sotiriou, Vincent Detours

## Abstract

A-to-I editing substitutes inosines for adenosines at specific positions in mRNAs and can substantially alter a cell’s transcriptome. Currently, little is known about how RNA editing operates in cancer. Transcriptome analysis of 68 normal and cancerous breast tissues revealed that the editing enzyme ADAR acts uniformly, on the same loci, across tissues. Controlled ADAR expression experiments demonstrated that the editing frequency at all loci is proportional to both ADAR expression levels and the individual locus’ editability—a propensity to be edited determined by the surrounding nucleotide sequence. Comparison of tumor transcriptomes to those of normal breast and breast organoids, i.e. pure normal breast epithelial cells, demonstrated that the editing frequency is increased in tumor cells. This was consistent with ADAR immunohistochemistry. We also demonstrated that type I interferon response and ADAR DNA copy number explain together 53% of ADAR expression in breast cancers, an observation also valid in nearly all of 20 other cancer types in The Cancer Genome Atlas. Interferon exposure increased ADAR mRNA, protein expression and editing in four breast cell lines. Finally, ADAR silencing using shRNA lentivirus transduction in breast cancer cell lines led to more cell proliferation and less apoptosis. Our results reveal that A-to-I editing is a pervasive, yet reproducible, source of variation that is controlled by two factors, 1q amplification and inflammation, both highly prevalent among human cancers. This suggests the potential for a new class of therapeutic targets and an unexpected role for inflammation in cancers.

## INTRODUCTION

While intense effort is currently being dedicated to cancer genome sequencing, comparatively little attention has been devoted at understanding how faithful RNA sequences are to the DNA sequences from which they were derived. Messenger RNA (mRNA) is the target of a series of post-transcriptional modifications that can affect its structure and stability, one of the most relevant being RNA editing *(1–3)*. The most common form of RNA editing in humans, the A-to-I type, is catalyzed by the <u>a</u>denosine <u>d</u>eaminases that <u>a</u>ct on <u>R</u>NA (ADARs) family of enzymes, which bind double stranded RNA (dsRNA) and turn adenosines into inosines at precise positions *(1, 3)*. Inosines are subsequently interpreted as guanosines by the cellular transcription machinery. ADAR enzymes are essential in mammals *(4, 5)* and exist in three forms: ADAR (also known as ADAR1), which is ubiquitous and has two isoforms—p110 is constitutive and p150 is inducible; ADARB1 (also known as ADAR2), principally expressed in the brain; and ADARB2 (also known as ADAR3), which contrary to ADAR and ADARB1 seems to be enzymatically inactive *(6, 7)*.

A-to-I edits can profoundly influence cellular functions and regulations by altering mRNA splicing, localization and translation, and interfering with the binding of regulatory RNAs *(8, 9)*. In addition to mRNA, ADAR can target non-coding RNAs such as micro RNAs (miRNAs), small-interfering RNAs (siRNAs) and long non-coding RNAs (lncRNAs), affecting both their structure and activities *(10–13)*. A-to-I editing has been shown to occur predominantly in highly repetitive *Alu* sequences, likely because their frequency (>10^6^) in the human genome make their arrangement in quasi-palindrome configurations prone to RNA duplex formation highly probable *(2, 9, 14, 15)*. High-throughput sequencing studies suggest that tens of thousands of positions are targeted by A-to-I editing in the human transcriptome *(16–19)*, and recent investigations report that potentially all adenosines in specific *Alu* repeats undergo A-to-I editing *(15)*.

Currently, a limited number of studies on A-to-I RNA editing in cancer have been published with the findings pointing to a diversity of effects. For example, in brain cancer editing inhibits cell growth and is reduced in glioma *(20, 21)* and pediatric astrocytoma *(22)*. In contrast, A-to-I editing increases during chronic myeloid leukemia progression *(23)*. In hepatocellular carcinoma, A-to-I editing of the antizyme inhibitor 1 (AZIN1) increases and neutralizes a key inhibitor of the polyamine synthesis pathway, thereby promoting proliferation *in vitro* and increasing tumor initiation and volume in a mouse xenograft model *(24)*. The studies published so far included a small number of tumor samples—an important limit given the sheer diversity of transcriptomes—and/or investigated a limited number of editing sites. Whether the edited transcript originated from cancer cells or other cell types, e.g. immune cells, present in the tumor mass was not addressed. Hence, both the magnitude and mechanisms regulating A-to-I editing in the majority of cancers, including breast cancer (BC), remain largely unknown.

The main objective of this study was to investigate and characterize the extent of A-to-I RNA editing in BC and define the principles governing the editing process in this as well as other types of cancer. We demonstrate that A-to-I edited mRNA loci are conserved both in matched normal and tumor breast tissues and across different patients. The editing frequency is, however, markedly increased at all editable loci in tumor compared to normal tissues. The number of edited loci and their editing frequency were highly correlated with ADAR expression. In breast and the majority of other cancers, ADAR expression appears to be regulated by two main factors: the type 1 interferon response and *ADAR* copy gains. Furthermore, ADAR silencing using shRNA lentivirus transduction in breast cancer cell lines led to a decrease in cell proliferation and an increase in apoptosis. Our results show that A-to-I editing is a pervasive phenomenon in cancer that can drive aberrant transcriptome expression associated with a pro-proliferative and an anti-apoptotic phenotype in breast cancer suggesting its potential role during carcinogenesis.

## METHODS

The methods are fully detailed in the Supplementary Methods available online. Briefly, the exome and transcriptome of 58 well characterized BC samples representing the four main known subtypes based on immunohistochemistry, namely TN, HER2+, luminal A and luminal B, and 10 matched normal samples were profiled using exome sequencing and RNA-Seq in paired-end mode on the Illumina HiSeq 2000 platform. Gene expression and SNPs profiles were obtained with Affymetrix^®^ HG-U133 Plus 2.0 Array chips and Affymetrix^®^ Genome-Wide Human SNP Arrays 6.0 for 57 and 49 tumor samples, respectively. RNA reads obtained from RNA-Seq were aligned simultaneously on the human genome and all known exonic junctions. Variant calls were submitted to a series of filters limiting artifact associated with RNA-seq. The identified RNA-DNA differences (RDDs) were validated in an independent cohort of 15 BC samples; moreover, few events as well as their edit frequency were validated using an independent technology (Roche FLX sequencer). The effect of interferon (IFN) on ADAR expression and editing was evaluated on 6 BC cell lines and one immortalized, non-transformed mammary epithelial cell line, MCF-10A. Cell lines were treated for 1, 2 or 5 days with IFN α, β, or γ. The effect of treatment on ADAR p110 and p150 protein and gene expression levels were evaluated quantifying the immunoblot signals and qRT-PCR, data respectively, while the effect of IFN treatment on editing distribution and frequency was investigated using amplicon sequencing (Roche FLX sequencer). In each sample, the mean editing frequency was correlated with clinico-pathological parameters and the expression of ADAR. The intracellular localization of ADAR was defined using IHC. The association between editing and *ADAR* amplification and/or a surrogate of interferon response (STAT1 expression) was evaluated in breast and 19 additional cancer types obtained from TCGA. Experimental assays and computational analyses are detailed in the Supplementary Methods.

## RESULTS

### Detection and validation of A-to-I editing sites in breast tissue

The extent of A-to-I RNA editing in BC was investigated by paired exome and transcriptome sequencing of a broad series of BC samples representing the major intrinsic subtypes including 17 triple negative (TN), 14 HER2-positive (HER2), 16 luminal A (LA) and 11 luminal B (LB) tumors (Supplementary Table S1). Paired exome and transcriptome sequencing of matched, tumor-adjacent normal tissue was performed on ten cases from this series. RNA-DNA single nucleotide differences (RDDs) were called as outlined in Supplementary Fig. S1 (details in Supplementary Methods).

Overall, we detected 16,027 RDDs in one or more samples, with all possible base changes represented (Fig. 1A). Among these, 560 RDDs were located in *Alu* regions and all were of the A-to-I type (Fig. 1A and Supplementary Table S2), consistent with the notion that A-to-I editing occurs predominantly in forward-facing *Alu* forming dsRNA duplexes processed by ADAR. Forty seven percent of the A-to-I *Alu* RDDs were present in the DARNED RNA editing site database *(25)*. In contrast, only 2.5% of A-to-I, non-*Alu* RDDs and 0.6% of non A-to-I RDDs were found in the DARNED database (Fig. 1B).

**Figure 1.**
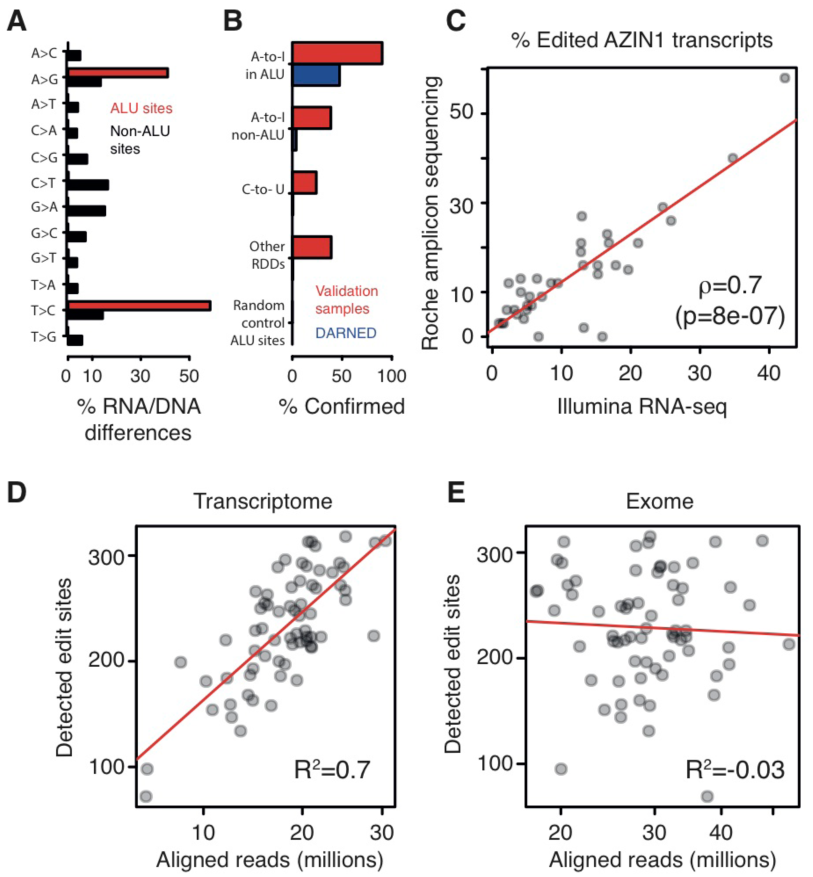
Detection of A-to-I editing. **(A)**, Substitution frequencies of RDDs. **(B)**, Percentage of RDDs confirmed in the validation dataset, N=15 BCs (in red), and the DARNED database (in blue). The negative control set is composed of 1,000 sites selected at random positions in randomly selected *Alu*s. **(C)**, Each dot represents a sample for which the frequency of edited AZIN1 transcripts has been measured with Illumina full transcriptome sequencing (x-axis) and Roche FLX amplicon sequencing (y-axis). ρ denotes the Spearman’s correlation. **(D)**, **(E)**, Number of detected *Alu* A-to-I sites as a function of transcriptome and exome coverages.

Breast tissue is not well represented in the studies covered by the DARNED database. Since gene expression and RNA editing frequency (defined for each sample as the ratio of the number of RNA-seq reads documenting the non-reference base relative to the total number of reads covering the site) could be regulated in a tissue specific manner, we further validated our findings on an independent breast series. This independent validation series included 15 BC samples with paired transcriptome and full genome sequencing data from the Sanger Institute. The genomic coordinates of our putative RDDs and the coordinates of 1,000 random *Alu* positions were sent to the Sanger Institute without any additional information. This blind test confirmed 90% of the *Alu* RDDs, while only one of the 1,000 random *Alu* sites was detected in the validation series. Beyond *Alu*, overlap with the validation series was below 40% (Fig. 1B). Given the low confirmation rate of RDDs located outside of *Alu* regions in both the DARNED database and the independent validation series and that the majority of human editing events are A-to-I detected in *Alu* repeats *(2, 9, 14, 15)*, our subsequent analyses focused exclusively on the subset of A-to-I RDDs located in *Alu* sequences. Because antizyme Inhibitor 1 (*AZIN1*) has also repeatedly been shown to be edited, this target was also included in our analyses *(16–18, 24, 26–29)*.

To evaluate the accuracy of edited transcript frequencies measured in our full transcriptome data, we generated amplicons of the *AZIN1* editing site region for 36 samples that were then analyzed by an independent sequencing technology (Roche FLX sequencer). The edit frequencies measured from full transcriptome and amplicon sequencing were remarkably consistent (Fig. 1C and Supplementary Tables S2 and S3) and thereby validated the accuracy of these estimations.

Sequencing depth is a key factor in detecting single nucleotide variations *(15)*, leading us to ask whether exome and RNA sequencing depths could influence the number of detectable *Alu* edit sites. As anticipated, while the number was not dependent on exome sequencing coverage, it did greatly increase with transcriptome coverage (Fig. 1D and E; Supplementary Table S4). A plateau was not reached in our dataset, which had a maximum coverage of ~3×10^7^ reads/sample. This suggests that with higher transcriptome coverage additional A-to-I editing sites should be detectable in the breast transcriptome.

### More A-to-I editing was found in tumor compared to normal matched breast tissue

To determine whether A-to-I editing is specifically altered in BC, the mean editing frequencies across all edited sites were compared between matched normal and breast tissues for ten cases where paired exome and transcriptome sequencing data was available for the normal tissue. *AZIN1* has repeatedly been reported to be edited with a high editing frequency for *AZIN1* shown to promote carcinogenesis in hepatocellular cancer *(24)*. Therefore, we compared the specific edit frequency of the *AZIN1* transcript determined by high-depth amplicon sequencing (Roche FLX sequencer) between tumor and matched normal breast tissues. The global mean editing frequency and the *AZIN1* specific editing frequency were higher in tumor compared to matched-normal breast tissues (Fig. 2A and B).

**Figure 2.**
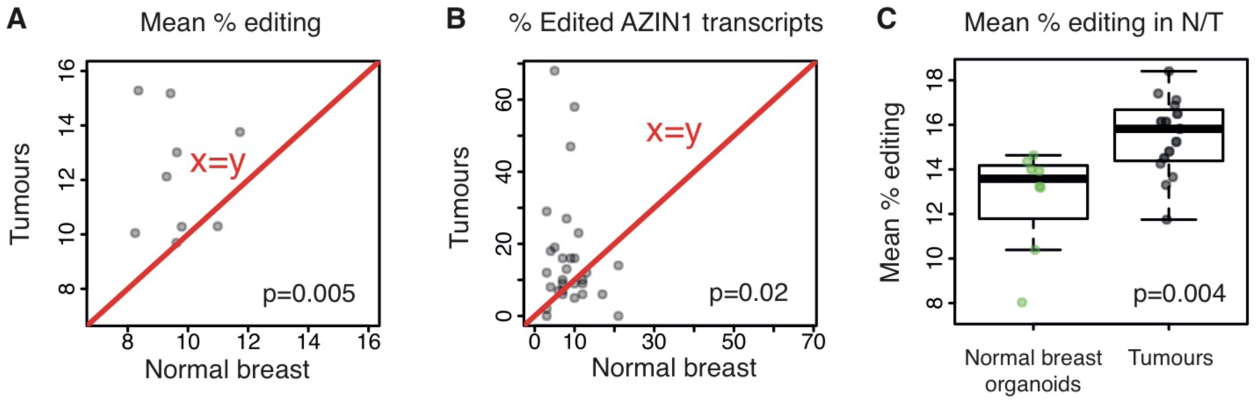
A-to-I editing and ADAR expression in normal and tumor breast tissue. **(A)**, Each dot represents a patient with the mean editing frequency in her normal (*x*-axis) and her matched tumor breast tissue (*y*-axis). **(B)**, Same as **(A)**, except that the AZIN1 editing frequency measured by Roche FLX amplicon sequencing is depicted. **(C)**, The mean editing frequency of 8 breast organoid cultures is compared to that of 15 breast tumors.

Lower epithelial cell content in normal compared to tumor breast tissue is a potential explanation for these observations. Thus, the site-averaged editing frequencies across all 560 *Alu* sites from the independent validation series (15 BCs) were compared to eight normal breast organoids (i.e. freshly isolated uncultured intact breast milk ducts). Editing was higher in tumor compared to normal cells (Fig. 2C), which validates our findings.

### Global A-to-I editing is governed by simple principles

The general principles governing A-to-I editing in BC were investigated in multiple, matched exome-transcriptome data pairs. The ADAR family of enzymes catalyzes A-to-I editing, leading us to first determine their expression levels in normal and tumor breast tissues as well as their association with editing frequency using transcriptome sequencing data. ADAR was expressed 9-fold more than ADARB1 and >1,000-fold more than ADARB2 (p<10^-16^, Supplementary Fig. S2A), which was anticipated because these last two isoforms are principally expressed in the brain. Moreover, while ADAR expression was higher in tumor compared to patient-matched normal breast tissues (p=0.005, Supplementary Fig. S2B), an inverse borderline-significant trend was observed for ADARB1 (p=0.1, Supplementary Fig. S2C).

The mean editing frequency (defined as the average editing frequency of all 560 *Alu* sites) was significantly positively correlated with ADAR mRNA expression levels (Fig. 3A; Supplementary Table S4), while it was weakly anti-correlated with ADARB1 expression levels (Supplementary Fig. S2D), as previously reported *(24)*. The global association detected between ADAR mRNA expression and the mean editing frequency was also observed at individual editing sites (Supplementary Fig. S3 and Supplementary Table S2). Considering both the high levels of ADAR mRNA expression and its strong correlation with the mean editing frequency found, our further analyses were focused on ADAR.

**Figure 3.**
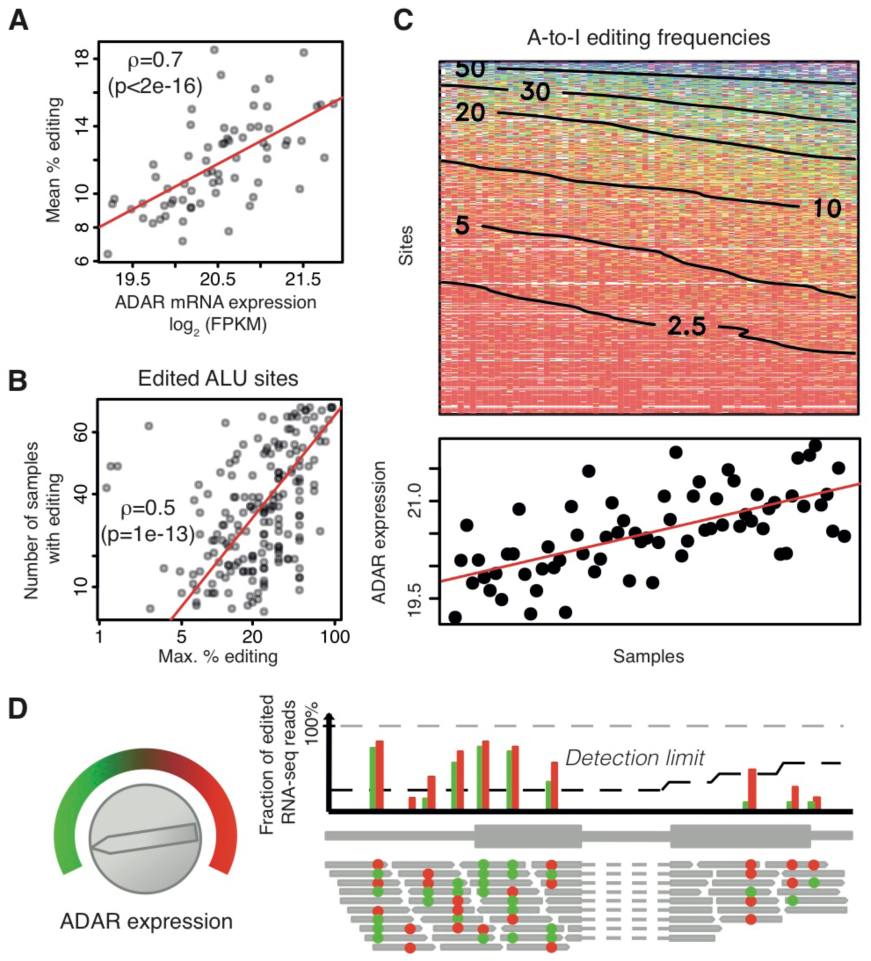
Model of A-to-I editing. **(A)**, Each dot represents a sample with its RNA-seq-estimated ADAR expression on the *x*-axis (in log_2_ of Fragments Per Kilobase per Million mapped reads), and its mean editing frequency across all 560 *Alu* sites on the *y*-axis. The RNA-seq expression of ADAR is highly correlated with microarrays and qRT-PCR expression (Supplementary Fig. S3). **(B)**, Each dot represents an *Alu* A-to-I editing site with the maximal edit frequency across all samples on the *x*-axis and the number of samples in which it was detectably edited on the *y*-axis. **(C)**, Heatmap of editing frequencies across all *Alu* A-to-I edit sites in all samples. Both are ordered by increasing (down-to-up, left-to-right) mean editing frequencies. Smoothed contour lines labels give the percentage of edited transcripts. The bottom panel shows corresponding ADAR expression. Negative controls are presented in Supplementary Fig. S3. **(D)**, Model of A-to-I editing. Turning the ADAR ‘expression knob’ clockwise increases ADAR expression. As a result, more transcripts are edited (red dots) and the editing frequency of all editable sites increases accordingly (compare green vs. red bars). Moreover, the detection limit at some sites for which editing was previously undetectable was passed. The detection limit depends on sequencing coverage, which is lower on the right-most exon. Importantly, the ranking of editing frequencies of the different sites is unaltered by *ADAR* expression.

Editing site distribution across normal and BC tissues was investigated by plotting the maximum edit frequency for all editing sites against the number of samples where editing of these sites was detected (Fig. 3B). These two variables were highly correlated indicating that if a site was highly edited in one sample it was very likely to be edited in many other samples. This also suggested that the editing sites are conserved between normal and tumor tissues and among BC patients.

The editing frequencies of all 560 A-to-I *Alu* editing sites were ordered by site and sample using an increasing average editing frequency (Fig. 3C; negative controls in Supplementary Fig. S3), which revealed that high editing frequencies were present in the samples with more editing sites and high ADAR expression. Conversely, samples with lower ADAR expression had fewer edited sites, which were edited at lower frequencies. Taken together, these data suggest a simple quantitative model of A-to-I editing (Fig. 3D). In this model, turning-up the ADAR expression ‘knob’ leads to detectable editing at more sites and a proportionally increased editing frequency of all the editable sites. Conversely, when ADAR expression is low, editing is detectable at fewer sites and at a lower frequency.

### Validation of the A-to-I editing model

We challenged this A-to-I editing model by inducing ADAR expression in four breast cell lines (three tumor and one normal tissue derived cell lines) with interferon alpha, a known ADAR inducer *(30)*. The effect of inducing ADAR overexpression on the editing frequency of *AZIN1* and four of the most edited *Alu* regions in the discovery series was analyzed by amplicon sequencing (Roche FLX sequencer). These experiments demonstrated: First, that the same sites were edited in all cell lines (Supplementary Fig. S4 and Supplementary Table S5), including 90 of the 91 sites detected by whole transcriptome sequencing *in vivo*. Second, the editing frequency profiles were conserved across all cell lines (Supplementary Fig. S4). Third, increasing ADAR induction time increased editing frequencies at all edited positions (Fig. 4A and Supplementary Fig. S4). Fourth, increasing ADAR induction time and/or the depth of coverage increased the number of detected editing sites (Fig 4B). Due to deeper coverage (typically >1,000X for the Roche FLX sequencer) of the cell line amplicons, we identified 137 new sites in addition to the 90 in the discovery dataset, which suggests there are likely even more sites to identify in breast tissue.

**Figure 4.**
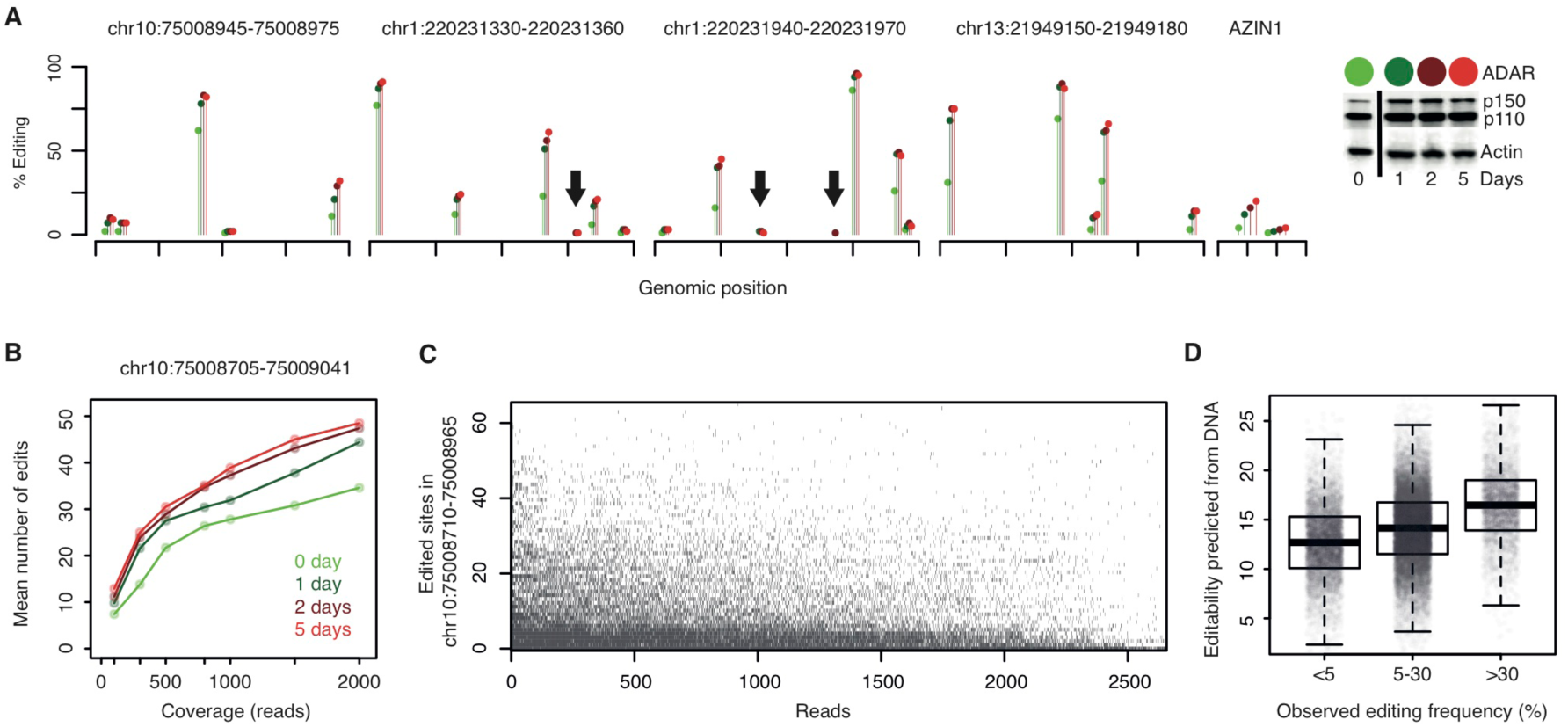
Validation of the A-to-I editing model. **(A)**, Effect of increasing ADAR expression in the cell line MCF7 on editing in 4 representative *Alu* regions and AZIN1. The full-length of sequenced regions are shown in Supplementary Fig. S4 for MCF7 and 3 more cell lines. Complete ADAR Western blots quantifications underlying the color scale are provided in Fig. 6E and in Supplementary Fig. S9A (see baseline t=0 and IFN-α, t ∊ {1, 2, 5} days tracks). Increasing ADAR expression increases the editing frequency at all editable positions, as predicted by the model of Fig. 3D. Similar results were obtained for IFN-β and IFN-γ (global, position-less, view Fig. 6F). Arrows point at editing sites detectable only at higher ADAR expression in our assay. **(B)**, Increasing sequencing coverage (*x*-axis) or ADAR expression (color scale) increases the number of detectable editing sites (*y*-axis). Coverage variation was implemented by down-sampling the total pool of sequencing reads, starting from 2,000x, down to 100x, and re-running the variant detection pipeline for each down-sampled alignment. Each data point is the mean of 10 down-sampling experiments. Error bars were omitted for the sake of clarity. **(C)**, Editing of individual mRNA molecules. Each black dot depicts an edited base (*y*axis) in a sequencing read (*x*-axis) spanning the entire 256bp region. Reads and sites were ordered by decreasing editing frequencies. Hundred and eighty-five non-edited reads were omitted from the figure. **(D)**, Each dot represents an *Alu* A-to-I site with coverage >20x from high coverage sample GM12878. Measured editing frequencies, *x*-axis, are compared to the editability score computed from the DNA sequence alone, *y*-axis. The calculation was run on 25,681 sites not used to develop the model.

We took advantage of the long reads (>300bp) and high coverage in the Roche FLX data to further validate our model by applying it to thousands of individual mRNA molecules transcribed from the same DNA region in the same individual. Focusing on one 256bp *Alu* region in one cell line, 65 of 68 adenosines potentially targeted by ADAR (Fig. 4C) were edited in at least one of the 2,842 mRNA molecules analyzed. The number of edited positions per transcript was highly variable, ranging from 0 to 26 (38% of all adenosines). As expected, the sets of edited positions in “low-edited” mRNA molecules tended to be subsets of those edited in “high-edited” mRNA molecules. These findings further validate our A-to-I editing model. Nevertheless, the editing process had a strong stochastic component at the level of individual molecules. Interpreting editability as a probability of edition by ADAR reconciles the molecule-level randomness with the predictable editability observed at the level of transcripts populations.

### Modeling editability

Beyond a dependency on ADAR mRNA levels, increases in editing frequencies at any given position may be proportional to an intrinsic, sequence-related property called ‘editability’. Editability depends upon biophysical interactions between an individual site with its surrounding RNA sequence and partnering as a duplex with ADAR. Because editability is zero at the vast majority of positions, the activity of ADAR appears to be site-specific.

In principle, editability should be partially predictable from sequencing data, which led us to develop and validate a simple proof-of-principle statistical model for editability. The model relies on the notion that: first an edited site must be part of an RNA duplex (i.e. in a sequence with a nearby palindromic match) and second ADAR activity is dependent on a specific nucleotide sequence in the vicinity of the edited base (details in Supplementary Fig. S5 and Supplementary Methods). To build the model, we analyzed the edit frequencies of 51,621 edited *Alu* sites with ≥20X coverage from an independent sample sequenced at very high coverage *(26)*. These sites were then ordered by genomic position. Half were used to fit a statistical model of edit frequencies based on DNA data alone (Supplementary Fig. S5). Editability scores were then computed for the other half, revealing that they are strongly associated with the observed edit frequencies (Fig. 4D). Our validated statistical model supports the notion that the edit frequency of an individual site in a given sample is controlled both by a site-specific factor, its editability, and a global factor, ADAR expression.

### Association of ADAR expression, A-to-I editing and clinico-pathological variables

The relevance of ADAR expression to the A-to-I editing process led us to analyze its tissue and cellular localization by immunohistochemistry (IHC). Uniform ADAR expression was detected in cancer cells (Fig. 5A) but to a lesser extent in normal cells and tumor-infiltrating lymphocytes (TILs; see Fig. 5B). Moreover, ADAR staining was markedly stronger in nucleoli (Fig. 5C), in agreement with previous findings *(31, 32)*.

**Figure 5.**
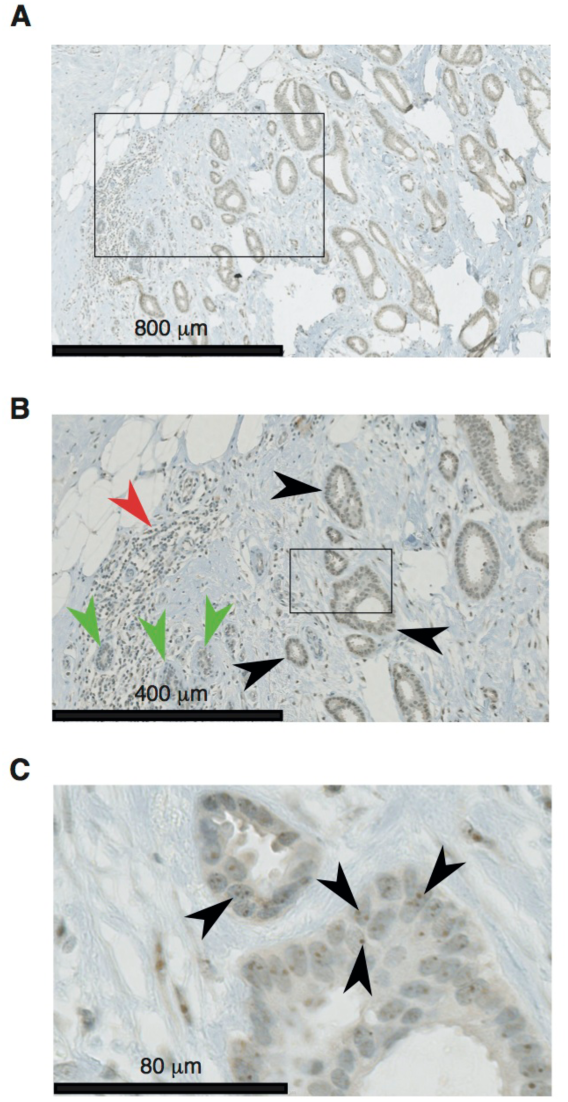
(**A**) Representative ADAR staining of a luminal A tumor. **(B)**, Zooming in panel **(A)** reveals that tumor staining (black arrows) is higher that in normal epithelium (green arrows) and lymphocytes (red arrows). **(C)**, Zooming further in panel **(B)** reveals a higher staining of nucleoli (black arrows).

To investigate the potential clinical impact of A-to-I editing, we determined whether the mean editing frequency was associated with tumor cell content (i.e. the proportion of malignant epithelial cells, adipose, stroma, normal epithelial cells and TILs) and/or well-established clinico-pathological parameters, including estrogen receptor, progesterone receptor, the proliferation marker Ki67, HER2 status, tumor size, nodal status and histological grade. The mean editing frequency was positively correlated with the percentage of TILs (Spearman’s correlation ρ=0.3, p=0.02), tumor size (ρ=0.3, p=0.01) and HER2 IHC staining (ρ=0.3, p=0.01; see Supplementary Fig. S6A-C and Supplementary Tables S1 and S4). Multivariate analysis of this dataset suggests that TILs and HER2 IHC are dependent variables in their association with editing frequency (Supplementary Fig. S6G).

To circumvent our limited sample size, correlations between these variables and ADAR expression were assessed in a large cohort of 787 BC patients with HER2 analyzed by IHC *(33)*. TILs were not scored in this series so the level of Signal Transducer and Activator of Transcription 1 (*STAT1*) expression, a proxy for type I interferon response, was used instead. This independent BC series confirmed an association between ADAR and STAT1 expression but not for HER2 status or tumor size (Supplementary Fig. S6D-F). The lack of an association with estrogen receptor, Ki67 and HER2 indicates that ADAR expression is not correlated with a specific BC subtype beyond their link with adaptive immune response.

### The interferon response and gains in ADAR copy number independently control A-to-I editing in cancer

The biological processes potentially associated with RNA editing were investigated by searching for genes whose expression had a strong positive correlation with the mean editing frequency (details in Supplementary Methods). Remarkably, 62 of the 85 genes identified were located on chromosome 1q (p=10^-66^). Since *ADAR* is located on chromosome 1q, we next used SNP data to determine *ADAR* copy numbers in our samples. *ADAR* amplification was frequent in our series (44%) and correlated with high mean edit frequencies (Fig. 6A).

**Figure 6.**
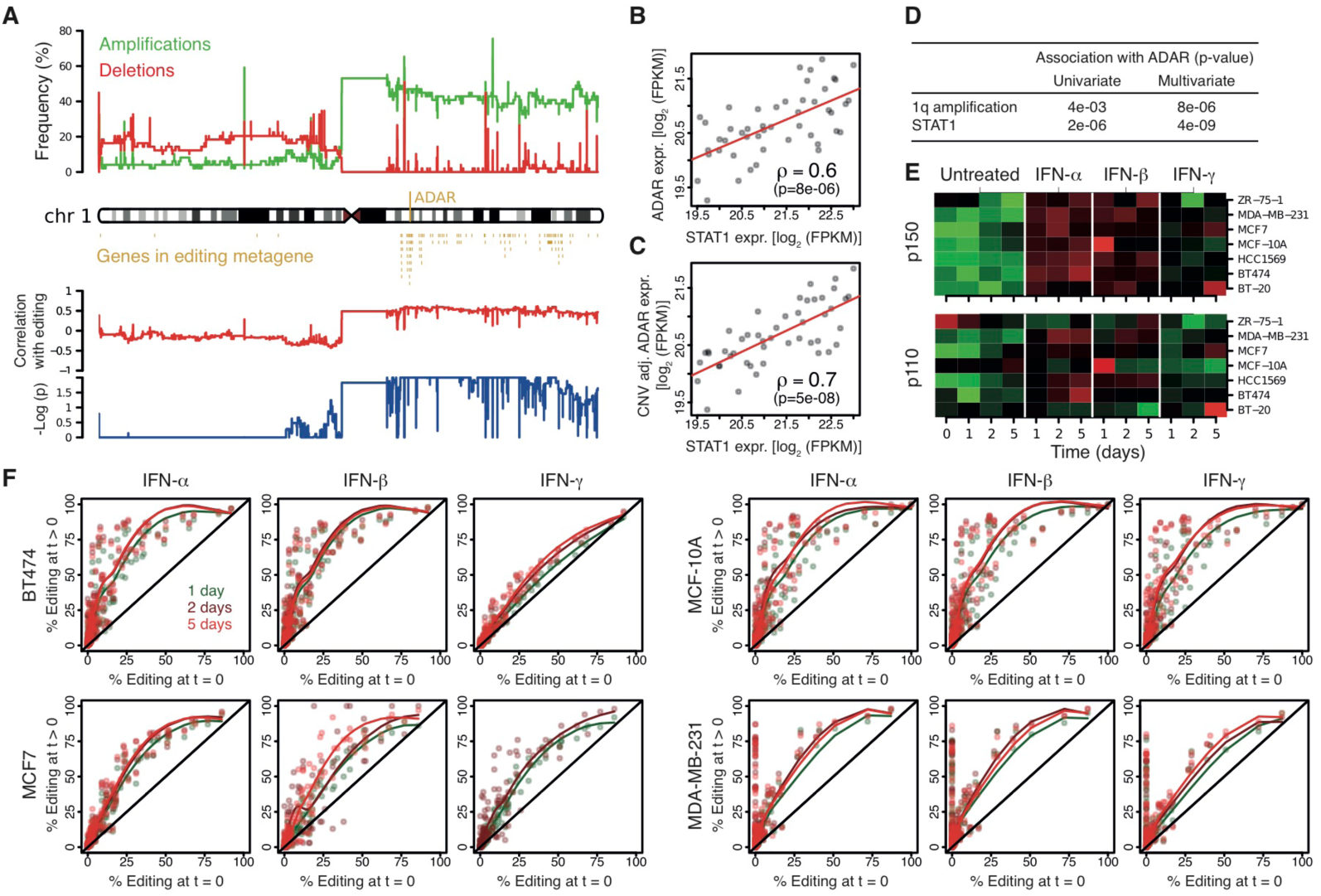
*ADAR* amplification and the interferon response are independent predictors of ADAR expression in cancer. **(A)**, The top panel shows the frequencies of amplifications/deletions along chromosome 1 in our series. The middle panel shows the genes whose expression is highly associated with that of ADAR. Nineteen genes not located on chr1 are omitted. The bottom panel shows the Spearman’s correlation coefficient and associated *p*-values of non-segmented copy-number array probes with the sample-wise mean editing frequencies. **(B)**, Dots represent tumor samples, with STAT1 expression on the *x*-axis and ADAR expression on the *y*-axis. **(C)**, Same as **(B)** with *ADAR* expression adjusted for *ADAR* copy number. **(D)**, Association p-values of *ADAR* copy number and STAT1 expression with ADAR expression increase in a multivariate analysis, demonstrating that ADAR expression is independently associated with these two variables. **(E)**, Seven breast cancer cell lines were exposed to interferon α, β and γ for 1, 2 and 5 days. Western blots quantifications are depicted for each cell line, interferon and time. Because expression dynamic ranges vary among cell lines, each line has its own color scale extending from low expression in green to high expression in red. The underlying gels are presented in Supplementary Fig. S9A and blot quantification in Supplementary Table S6. **(F)**, Editing frequencies in the absence of treatment (*x*-axis) vs. interferon treatment (*y*-axis). Points depict the editing sites in AZIN1 and the 4 *Alu* regions of Supplementary Fig. S4. Points are above the identity line *x*=*y* (black diagonals), i.e. interferon increase editing frequencies at all sites. Library preparation failed for MCF7/IFN-γ at 5 days. Limited sequencing coverage precluded detection of some editing events for MDA-MB-231, t=0 and t=1 days.

Chromosome 1q contains hundreds of genes and therefore its amplification could have a systemic impact on the BC transcriptome *(33)*. Therefore, we further characterized the genes correlated with editing that were independent from 1q amplification. First, the microarray expression data were adjusted for 1q copy number to remove any potential confounding effects of *ADAR* amplification and then gene set analysis was performed *(34)* to identify canonical pathways associated with the mean edit frequency. The 13 significant pathway gene sets revealed by this analysis were all involved in interferon responses, interferon-related DNA and RNA sensing and lymphocyte biology (Supplementary Fig. S7A). We also investigated gene sets with shared transcription factor binding motifs between their promoters. The 7 significant gene sets identified were overwhelmingly related to *NFκB* and the interferon response, including the Interferon Response Factors *IRF1*, *IRF2* and *IRF7* (Supplementary Fig. S7B). To further investigate the relationship between interferon-related genes and ADAR expression, the median expression levels of STAT1 (Fig. 6B) and 389 type I interferon-inducible genes (Supplementary Fig. S7C) derived from 10 microarray studies *(35)* were measured. The expression of STAT1 and the 389 genes were positively associated with ADAR expression, suggesting that increased editing was part of a broader type I interferon response related to the chronic inflammatory state in cancer.

The respective roles of *ADAR* copy number and STAT1 expression (as a proxy for interferon response) in the A-to-I editing process were further defined using multivariate analysis to demonstrate that they are independently associated with *ADAR* expression (Fig. 6B-D). When STAT1 was a strong predictor of ADAR expression (Fig. 6B), it could be strengthened by adjusting ADAR expression for *ADAR* DNA copy number (Fig. 6C). Taken together, STAT1 and ADAR copy number explained 53% of ADAR expression variation. The independent effect of type I interferon response and *ADAR* amplification was also supported by measuring the constitutive p110 and interferon-inducible p150 ADAR isoforms (Supplementary Fig. S8). STAT1 expression was more strongly correlated with p150 than p110 and conversely *ADAR* copy number more with p110 than p150.

While *ADAR* amplification is likely limited to malignant epithelial cells, the type-I interferon effect could be principally mediated by TILs. Seven breast cell lines (derived from the four principal BC molecular subtypes and normal breast) were treated with individual interferons (α, β, and γ) to determine if editing can be directly increased by interferon. ADAR p150 protein expression increased with all three interferons in all cell lines at each time point (Fig. 6E, Supplementary Fig. S9A), while p110 induction was weaker and less consistent. The moderate but significant correlation between p110 and STAT1 mRNA detected in primary tumors suggests that a small amount of p110 was induced (Supplementary Fig. S8F). The same four cell lines used to validate our A-to-I editing model were analyzed for p150 and p110 ADAR mRNA isoform expression levels, the editing proportion of AZIN1 and the four most edited *Alu* regions previously selected. The mRNA levels for p110 and p150 isoforms paralleled their protein expression (Supplementary Fig. S9B). Moreover, editing increased at all editable sites with all interferons in the four cell lines (Fig. 6F), although higher editing levels were observed at 2 or 5 days compared to untreated or 1 day. The induction of ADAR and editing was lowest for IFN-γ. These experiments confirm that type-I interferon response affect A-to-I mRNA editing in epithelial cells.

### ADAR is involved in cell proliferation and apoptosis in breast cancer

As we have shown that ADAR expression and mean editing frequency were higher in breast tumors compared to matched normal tissues, we aimed to further investigate its role on cell proliferation, migration and apoptosis. To that aim, ADAR expression was stably knocked down in 3 different representative breast cancer cell lines (MDA-MB-231, MCF7 and BT474) using shRNA lentiviral particles (shRNA ADAR). The 3 cell lines were transduced with scramble shRNA lentiviral particles (shRNA control) as a negative control for the functional experiments. ADAR silencing was confirmed by Western blot analysis (Fig. 7A).

**Figure 7.**
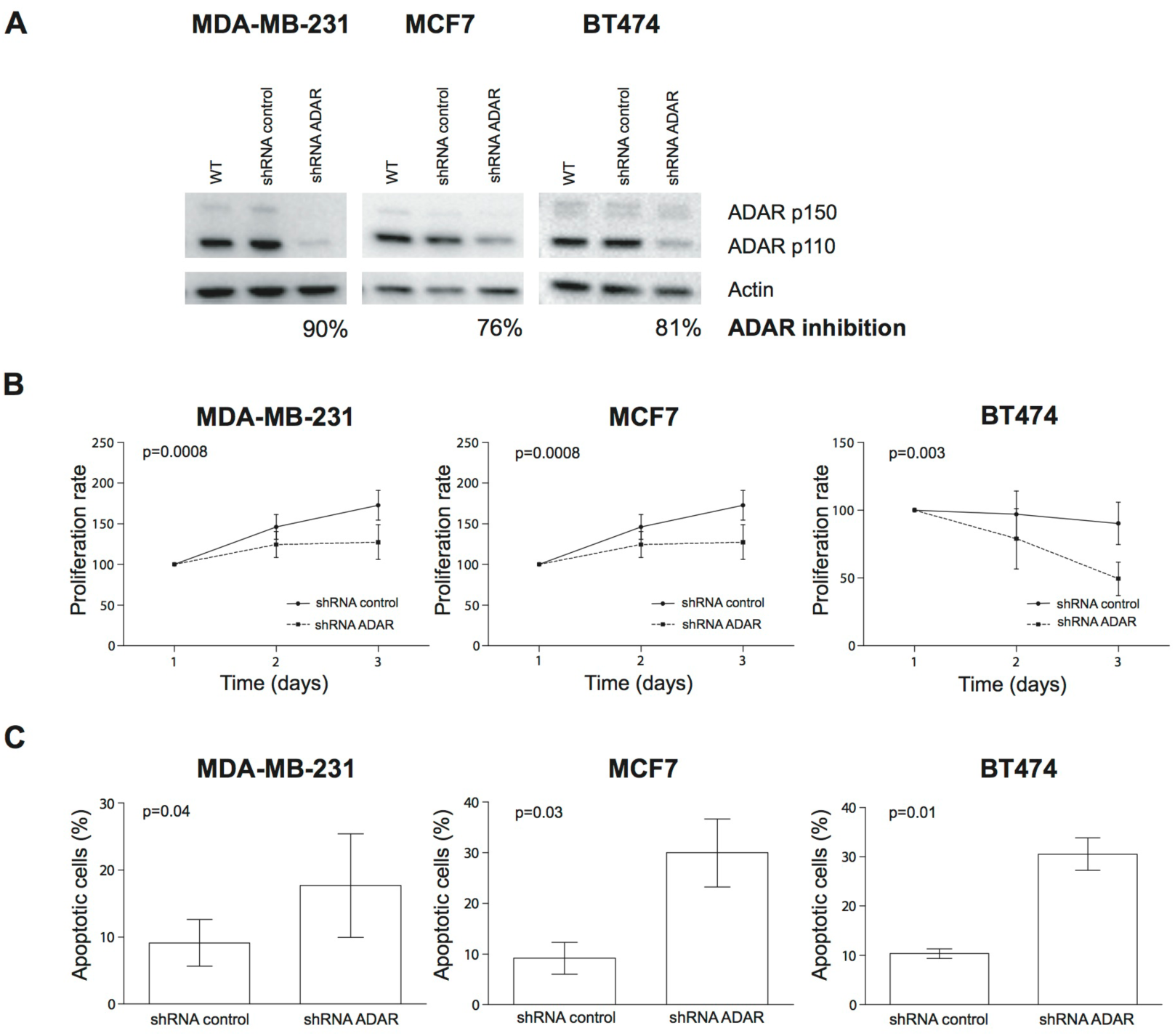
ADAR involvement in cell proliferation and apoptosis. **(A)**, Western blot analysis of ADAR silencing after shRNA lentiviral transduction in MDA-MB-231, MCF7 and BT474 breast cancer cell lines. **(B)**, ADAR silencing statistically decreases cell proliferation. Cell growth curves for ADAR-knock down cells (shRNA ADAR) and control cells (shRNA control) in MDA-MB-231, MCF7 and BT474 BC cell lines. **(C)**, ADAR silencing statistically increases cell apoptosis. Illustration of the percentage of apoptotic cells in ADAR-knock down cells (shRNA ADAR) and control cells (shRNA control) in MDA-MB-231, MCF7 and BT474 BC cell lines. Error bars depict standard deviation of three independent experiments.

To assess the role of ADAR in cell proliferation, MTT assays were performed. These experiments showed that ADAR silencing led to a statistically significant decrease in cell proliferation (shRNA ADAR) compared to the control cells (shRNA control) in all cell lines (Fig. 7B). These results suggest that ADAR promotes cell proliferation. No significant effect of ADAR silencing was found regarding cell migration. The role of ADAR in apoptosis was investigated using Annexin V assays. ADAR silencing led to a statistically significant increase in cell apoptosis (shRNA ADAR) compared to the control cells (shRNA control) in all cell lines (Fig. 7C) suggesting that ADAR may act as an anti-apoptotic factor.

### The role of ADAR gains and interferon responses in other cancers

*ADAR* amplification is frequent in human cancers (Fig.8) and inflammatory responses are pervasive in this disease. This information led us to investigate whether these two factors were related to ADAR expression in 4,480 cancers from the The Cancer Genome Atlas (TCGA) *(36)*, where sample-matched expression and copy number profiles are available. The representative analyses shown in Fig. 6B and 6C were reproducible across the TCGA dataset, which spans 20 types of cancer from 16 organs (Fig. 7). Overall, ADAR expression was consistently associated with both *ADAR* copy number and STAT1 expression. Similar to BC, adjusting ADAR expression for *ADAR* copy number increased the correlation between ADAR and STAT1 for all except pancreatic, kidney and thyroid tumors. The frequency of *ADAR* amplification was low in kidney and thyroid tumors, therefore correcting for *ADAR* copy number had a limited effect. These data suggest that ADAR expression could be principally driven by interferon in these two types of cancer. In most cancers, however, the editing process is driven by both type I interferon and *ADAR* copy number amplification. A correlation between ADAR copy number and ADAR expression has also been recently reported in esophageal cancer *(29)*.

**Figure 8.**
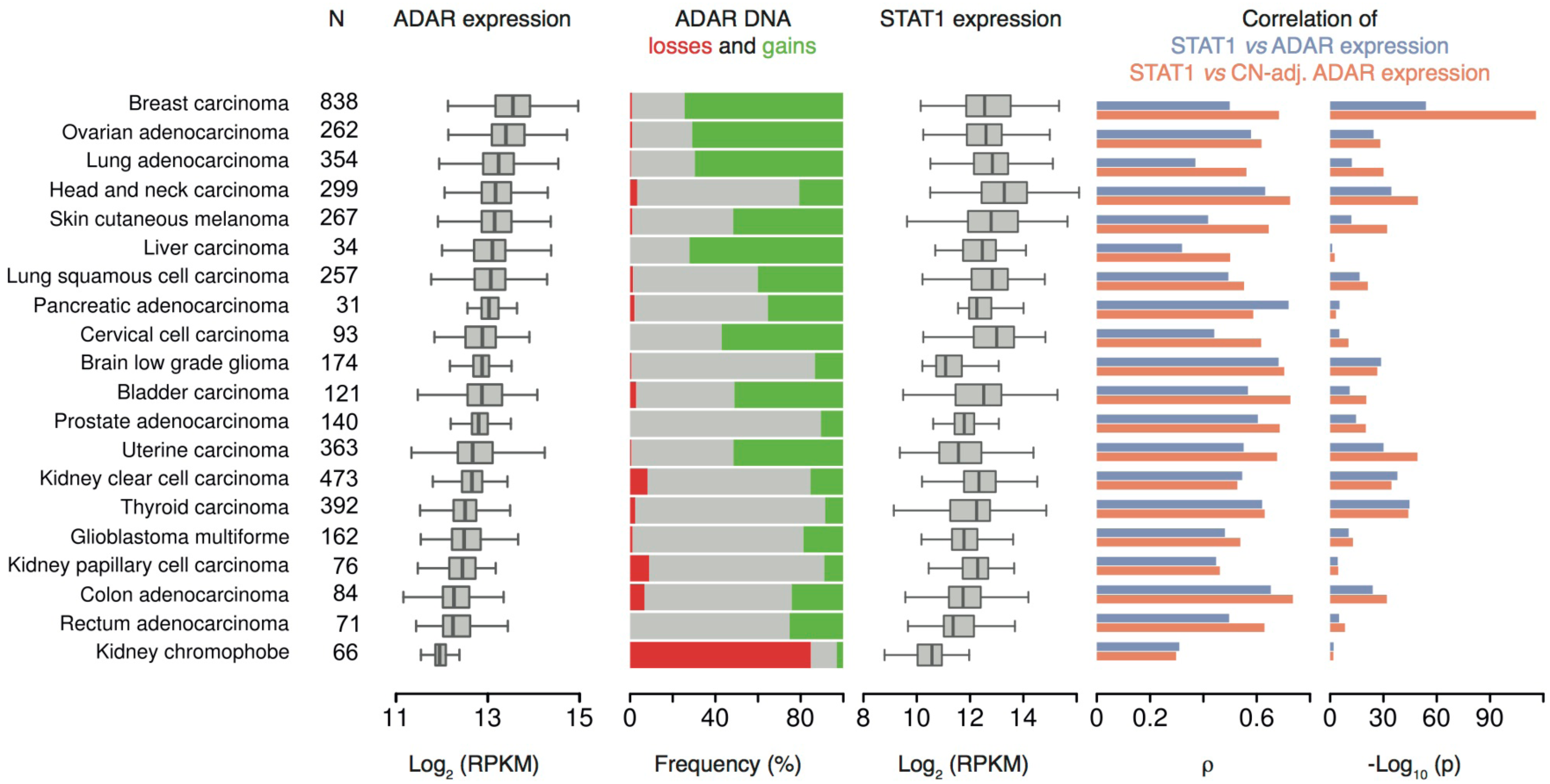
*ADAR* amplification and the interferon response predict ADAR expression in human cancers. We included all TCGA datasets and tumors (see ‘N’ column) for which both copy number and RNA-seq expression data (pipeline v3) were available. Datasets are ordered by decreasing median ADAR expression (top to bottom). The three leftmost plots depict the distributions of ADAR expression, *ADAR* DNA copy number and STAT1 expression across each dataset. The two rightmost bar plots extend to TCGA data the calculation presented for our data in Fig. 6B and 6C. In most cancers, adjusting ADAR expression for *ADAR* copy number increases the Spearman correlation, ρ, with STAT1 (compare the orange bars to the blue bars).

## DISCUSSION

Next generation sequencing has fostered greater interest in RNA editing as a mechanism that governs additional and unexpected layers of genomic complexity. Recent studies have focused on defining the optimal bioinformatic strategies for identifying reliable RNA editing sites *(16, 18, 26, 37, 38)*. Studies of the mammalian genome and transcriptome have shown that A-to-I editing, catalyzed by the ADAR family of enzymes, is the main form of RNA editing *(1–3, 15)*. At first this phenomenon appeared to be restricted to a few targets located in coding regions; however, it is now clear that most A-to-I editing sites are located in non-coding regions. In particular, these sites tend to cluster in *Alu* regions, which due to their highly repetitive nature, are prone to form double-stranded RNA structures recognized by ADAR family enzymes *(2, 9, 14, 15)*.

Initial studies found that the altered editing of specific targets were a determinant of tumor behavior, spawning efforts to define a potential role for A-to-I RNA editing in cancer initiation and development *(20, 39, 40)*. Solid and hematologic cancers have been investigated with a few targets identified *(24, 29)*; however, most of these studies performed an in depth analysis on small cohorts of patients. The magnitude of A-to-I editing in cancer as well as the mechanisms controlling and regulating the A-to-I editing machinery are currently unknown.

To address both points, we preformed the largest survey on RNA editing in cancer to date by profiling dozens of BCs with their matched healthy breast tissue. This was accomplished by using exome and transcriptome sequencing together with SNPs and gene expression arrays in combination with state-of-the-art bioinformatics analyses. The increased sample size of this study opened a window to principles governing A-to-I editing that were previously out of reach. A significant finding from our study is the demonstration that the same sites are edited in normal and tumor breast tissues as well as in several BC cell lines. We further show that while the editing frequency profiles are conserved across tissues and BC cell lines, the frequency of editing was significantly higher in tumors compared to their matched normal breast tissues. High editing frequencies were also detected in samples with more detectable editing sites and/or where ADAR expression was high. These data provided the basis for our A-to-I editing model, where increases in ADAR expression acts on all editable positions in the transcriptome proportionally to their editability. This model was successfully validated on BC cell lines. ADAR appears to have site-specific activity, which is partly influenced by the biophysics of interactions between nucleotides in the surrounding RNA sequences and their duplex partnering with ADAR, a property that we define as editability.

Longer ADAR induction times and/or deeper sequencing coverage increased the number of editing sites detected. Interestingly, no plateau was reached at the depths we investigated, with up to 3×10^7^ aligned reads per sample. A previous study made a similar observation using coverage up to 5×10^8^ mRNA reads/sample, where no plateau was reached despite >140,000 A-to-I sites detected in the *Alu*’s (26). Differences in number of edited sites between the cited work and the present study could also be due to the cell type analyzed (a lymphoblastoid cell line vs. breast tissue/cell lines, respectively) and the DNA sequencing strategies (whole genome vs. coding sequences and their neighborhood). It is anticipated that a large number of additional A-to-I editing sites beyond those identified here remain to be discovered in BC. The data presented here clearly demonstrate that A-to-I editing is a pervasive phenomenon in cancer, and suggest that it is a major source of mRNA sequence variability in breast and potentially other types of cancer. The importance of this observation is that editing has the potential to significantly impact transcriptional regulation and cellular functions in tumor cells. Indeed, our in vitro studies have shown that ADAR silencing decreases cell proliferation and promotes apoptosis supporting the potential carcinogenic role of ADAR and consequently A-to-I editing in breast cancer.

This study is the first to identify two key factors independently determining ADAR expression and thereby A-to-I RNA editing in breast and other human cancers: first the type-I interferon response in tumors, and second *ADAR* copy number gains. The mean editing frequency was found to be significantly correlated with the expression of STAT1 and type I interferon target genes, both in our patient series and a large pool of BC datasets. This observation suggests that the immune response profoundly impacts the transcriptome sequence of tumor cells and thereby influences the internal mechanisms that govern their behavior.

Recent work has revealed that aberrant expression of ADAR and APOBEC family proteins occurs in many human diseases, including cancer. In breast and other tumor types, mutational signatures are associated with APOBEC family proteins *(41)* with evidence that APOBEC-mediated mutagenesis is highly active in human cancers *(42, 43)*. Although, the role of ADARs and RNA editing in human disease is just beginning to emerge *(44)*, the link between A-to-I editing by ADAR and the type I interferon response shown here suggests that editing can not only affect tumor cell activities, but also those of immune and other cells present in the tumor microenvironment. A significant role for ADAR is further supported by our demonstration that its expression is significantly upregulated by *ADAR* copy number gains in breast (up to 75%) and other cancers (up to 70%). Overall, these data highlight the potential magnitude of A-to-I RNA editing in tumors and thereby the possibility for large-scale clinical consequences. RNA editing and/or APOBEC-mediated mutagenesis could shape the immunogenicity of the tumor and thereby directly affect anti- and/or protumor immune responses. RNA editing itself, the processes it regulates and its potential to differentially direct activities in response to the chronic inflammatory tumor microenvironment, may have important implications for clinical progression in breast and other cancers.

This study identifies the principles governing A-to-I editing in breast and potentially all types of cancer. These data further identify the type I interferon pathway as an important mechanism regulating this phenomenon. The wide-spread editing we observed, in combination with the conservation of editing sites detected across tissues and patients, suggests there are may be clinical and therapeutic implications for a wide range of cancer patients. However, modulating editing at an individual site is entangled with many processes. The model we established for A-to-I editing implies that modulation of ADAR will also affect all editable sites in expressed transcripts. In addition, ADAR has recently been shown to enhance miRNA processing *(45)*. Thus, variation in the editing frequency at an individual site is likely to be correlated with global variation of miRNA expression. Finally, variation of ADAR expression *in vivo* will possibly be associated with modification of the hundreds of genes located on 1q and/or controlled by interferon. Determining whether increasing A-to-I editing limits or enhances cancer progression will need to take into account all of these potential variables. More research is needed to identify the critical editing sites, establish their potential as markers of cancer evolution and investigate them as a new class of therapeutic targets.

## DATA ACCESS

Sequencing and SNPs array data obtained from the enrolled patients are archived at European Genome-phenome Archive, <u>https://www.ebi.ac.uk/ega</u>, under accession number EGAS00001000495; amplicon sequencing data obtained from cell lines are archived at European Nucleotide Archive, <u>http://www.ebi.ac.uk/ena/home</u>, under study accession number ERP004253; gene expression array data are archived at the Gene Expression Omnibus, <u>http://www.ncbi.nlm.nih.gov/geo</u>, under accession number GSE43358. Clinical information, results of sequence and arrays preprocessing and biological assays are available in the online Supplementary Tables.

## ACKNOWLEDGMENTS

We thank Raphael Leplae and the ULB Computing Center for their support, Roland De Wind for pathology support, Cédric Blanpain, Sabine Costagliola, Jacques E. Dumont, Pierre Vanderhaeghen and Gilbert Vassart for the helpful discussions.

